# FUN-LOV: Fungal LOV domains for optogenetic control of gene expression and flocculation in yeast

**DOI:** 10.1101/285296

**Authors:** Francisco Salinas, Vicente Rojas, Verónica Delgado, Javiera López, Eduardo Agosin, Luis F. Larrondo

## Abstract

Optogenetic switches permit accurate control of gene expression upon light stimulation. These synthetic switches have become a powerful tool for gene regulation, allowing modulation of customized phenotypes, overcoming the obstacles of chemical inducers and replacing their use by an inexpensive resource: light. In this work, we implemented FUN-LOV; an optogenetic switch based on the photon-regulated interaction of WC-1 and VVD, two LOV (Light Oxygen Voltage) blue-light photoreceptors from the fungus *Neurospora crassa*. When tested in yeast, FUN-LOV yields light-controlled gene expression with exquisite temporal resolution, and a broad dynamic range of over 1300-fold, as measured by a luciferase reporter. We also tested the FUN-LOV switch for heterologous protein expression in *Saccharomyces cerevisiae*, where Western blot analysis confirmed strong induction upon light stimulation, surpassing by 2.5 times the levels achieved with a classic *GAL4*/galactose chemical inducible system. Additionally, we utilized FUN-LOV to control the ability of yeast cells to flocculate. Light-controlled expression of the flocculin encoding gene *FLO1*, by the FUN-LOV switch, yielded Flocculation in Light (FIL), whereas the light-controlled expression of the co-repressor *TUP1* provided Flocculation in Darkness (FID). Overall, the results revealed the potential of the FUN-LOV optogenetic switch to control two biotechnologically relevant phenotypes such as heterologous protein expression and flocculation, paving the road for the engineering of new yeast strains for industrial applications. Importantly, FUN-LOV’ s ability to accurately manipulate gene expression, with a high-temporal dynamic range, can be exploited in the analysis of diverse biological processes in various organisms.

**Importance:** Optogenetic switches are molecular devices which allow the control of different cellular processes by light, such as gene expression, providing a versatile alternative to chemical inducers. Herein, we report a novel optogenetic switch (FUN-LOV) based on the LOV-domain interaction of two blue-light photoreceptors (WC-1 and VVD) from the fungus *N. crassa*. In yeast cells, FUN-LOV allowed tight regulation of gene expression, with low background in darkness and a highly dynamic and potent control by light. We used FUN-LOV to optogenetically manipulate, in yeast, two biotechnologically relevant phenotypes: heterologous protein expression and flocculation, resulting in strains with potential industrial applications. Importantly, FUN-LOV can be implemented in diverse biological platforms to orthogonally control a multitude of cellular processes.

## INTRODUCTION

Control of gene expression is an important tool for basic and applied research in biology. Chemical inducers, such as IPTG, methanol or galactose, have been profusely employed for inducible gene regulation despite their obvious limitations including potential interference with metabolic processes, difficult removal from the culture media once added, and insufficient temporal and spatial/dose resolution (1). In addition, the cost of chemical inducers can restrict their uses in some industrial applications, whereas temperature induction or constitutive gene expression are chosen based on cost efficiency regardless of their far-from-optimal characteristics (2). Light constitutes a promissory alternative for the control of gene expression, considering its low cost, reduced toxic effects, adjustable levels, and high temporal and spatial resolution (3). In several organisms, light readily controls different processes, including gene expression through photoreceptors with specialized domains which, under light stimulation, undergo a conformational change passing to an active state (4). Such photoreceptors have allowed defining basic building blocks from which to develop optogenetic switches, which can be engineered into synthetic light-controlled orthogonal transcription factors (5). During the past years, a nascent repertoire of optogenetic switches responding to light of different wavelengths, and assembled in different platforms, has become available(6-11).

Fungal photoreceptors have been an underexploited source of biological parts for the implementation of optogenetic switches. *Neurospora crassa*’s blue-light photoreceptor VIVID (VVD) has the capacity to self-dimerize upon light stimulation through its Light Oxygen and Voltage domain (LOV domain) (12). This feature of VVD already led to the development of the optogenetic system denominated “LightOn”, which was successfully utilized for light-controlled expression of transgenes in mice and mammalian cells (13, 14). Notably, in Neurospora, VVD also interacts with the blue-light photoreceptor White Collar 1 (WC-1), through WC-1’s LOV domain, allowing *N. crassa* to photoadapt in the presence of continuous illumination (15-17). This naturally-occurring LOV-LOV interaction of WC-1 and VVD opens the door for the development of novel optogenetic switches based on different LOV pairs, such as WC-1/WC-1 or WC-1/VVD.

The budding yeast *Saccharomyces cerevisiae* ranks amongst the most relevant and versatile microorganisms for biotechnology. The absence of photoreceptors in the yeast genome (18) has fuelled the implementation of different optogenetic switches in this biological chassis, allowing the orthogonal control of diverse cellular processes by light (19-21). However, despite their obvious advantages, the use of optogenetic switches to control biotechnologically relevant phenotypes in yeast has been seldom implemented (22). For example, optogenetic switches for heterologous protein expression in yeast would replace the addition of chemical inducers, reducing the cost of industrial scale bioprocesses, providing also a dynamic control of gene expression and the effective temporal control of the *on* and *off* states. Another biotechnological relevant operation in yeast fermentation is flocculation, which allows a fast, accessible and efficient way to remove remaining yeast cells after fermentation processes (23). Flocculation is controlled by the *FLO* genes, a family of subtelomeric genes, which trigger cell aggregation upon nutrient starvation or environmental stress conditions (24, 25). These features position flocculation as an ideal and biotechnologically relevant target for optogenetic control.

In this work, we present as proof of concept, a new optogenetic switch that provides an ample dynamic range of expression, with low-background in the off (dark) state. Its implementation in yeast provides evidence of tight regulation of gene expression by blue-light and the control of biotechnologically relevant features and phenotypes, such as heterologous protein expression and flocculation.

## RESULTS

### FUN-LOV provides a dynamic range of gene expression

The new optogenetic switch named FUN-LOV, was developed based on the pairing of WC-1 and VVD LOV domains, an interaction known to occur as part of the *N. crassa* photoadaptation process (17). The configuration of the FUN-LOV switch follows a Yeast Two-Hybrid system design logic, where the LOV domain of WC-1 is bound to a *Gal4* DNA binding domain (GAL4-DBD) and VVD’s LOV domain is tethered to the *Gal4* transactivation domain (GAL4-AD). Thus, upon the light stimulated interaction of this LOV domain pair, a functional TF is reconstituted, activating transcription as evidenced by a destabilized firefly luciferase reporter (*Luc*) under the control of the *GAL1* promoter (*P*_*GAL1*_) (Fig. 1A). Additionally, with the aim to increase even further gene expression induction, we designed a synthetic version of the *GAL1* promoter (*P*_*5xGAL1*_), which included four additional *GAL4-UAS* DNA binding site sequences (Fig. 1B).

**FIG 1.**
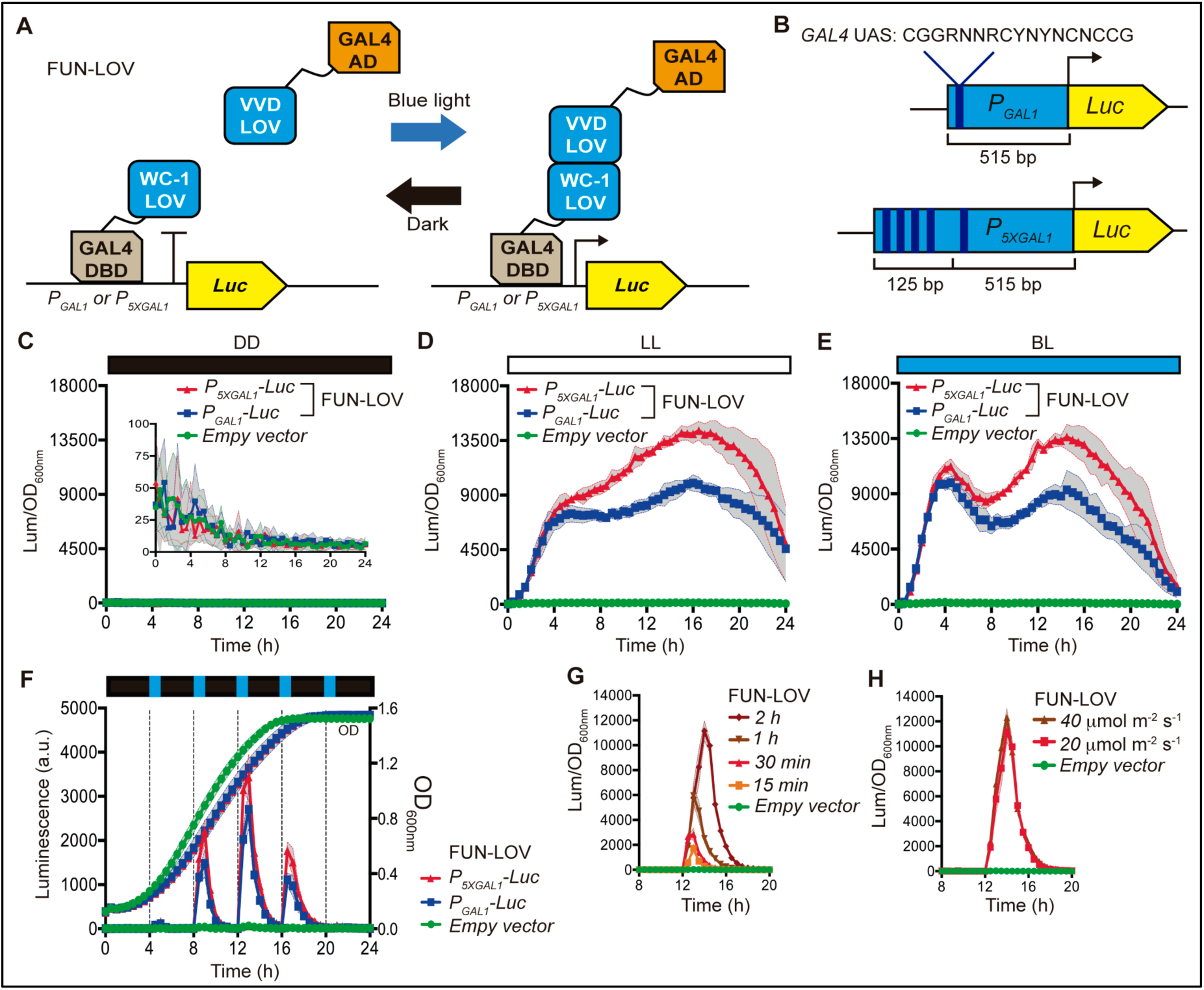
The FUN-LOV switch allows dynamic control of gene-expression by light. (A) FUN-LOV is based on a Yeast Two Hybrid design, where the LOV (**L**ight **O**xygen and **V**oltage) domains of WC-1 and VVD photoreceptors are tethered to the Gal4 DNA binding domain (DBD) and the Gal4 activation domain (AD), respectively. Upon light stimulation, the interaction between the LOV domains reconstructs a chimeric Gal4 transcription factor activating the expression of luciferase which is under the control of a *P*_*GAL1*_ or *P*_*5XGAL1*_ promoter. (B) Design of the synthetic promoter *P*_*5XGAL1*_, which contains four additional Gal4-UAS sequences upstream of the *GAL1* (*P*_*GAL1*_) promoter sequence. Luciferase expression controlled by FUN-LOV under constant darkness (DD) (C), constant white-light (LL) (D) and constant Blue-Light (BL) (E). (F) Temporal and dynamic expression of luciferase recorded in yeast cells grown for several hours and subjected to pulses of 30 min of Blue-light as indicated. (G) Blue-light pulses of increasing length, progressively augment luciferase expression. (H) Light intensities higher than 20 μmol m^-2^ s^-1^, applied for two hours, do not further increase luciferase levels. In (G) and (H), data corresponds to luciferase under the control of a *P*_*5XGAL1*_ promoter. Yeast growth was actively monitored at OD_600_, and directly plotted (F) or utilized to normalize luminescence data (Lum/OD_600_). In panels (C) to (H), each plot corresponds to the average of six biological replicas with its standard deviation represented as shadowed regions.

FUN-LOV revealed high levels of luciferase expression under constant white-light (LL) or constant blue-light (BL) conditions, with notably low levels of background expression in constant darkness (DD) (Fig, 1C, 1D, 1E; Fig. S1 and S2). In our hands, the expression levels achieved by the FUN-LOV switch were superior to those obtained applying classical galactose induction (Fig. S2). Furthermore, the maximum luciferase expression levels of the system were 1218-fold for white-light and 1316-fold of induction for blue-light, utilizing the synthetic *P*_*5xGAL1*_ promoter (Fig. 2A, 2B, 2C and 2D). The FUN-LOV system yielded similar results employing two different yeast genetic backgrounds (including the absence of endogenous Gal4/Gal80 proteins) and displayed robust luciferase expression either when maintained episomally or integrated in the genome (Fig. S1, S2 and FIG. 2). As luminescence was being measured directly from living cells, we wanted to confirm that luciferase levels from protein extracts would reflect the same range of induction. Therefore, luciferase activity was measured in extracts from cells grown under constant light (LL) or darkness (DD) for 8 hours, ratifying the strength of the FUN-LOV switch and revealing even higher levels of induction (∼2500-fold, Fig. 2E). Thus, the FUN-LOV switch provides remarkable high levels of gene expression, as measured by a luciferase reporter, under constant illumination with low background expression in darkness.

**FIG 2.**
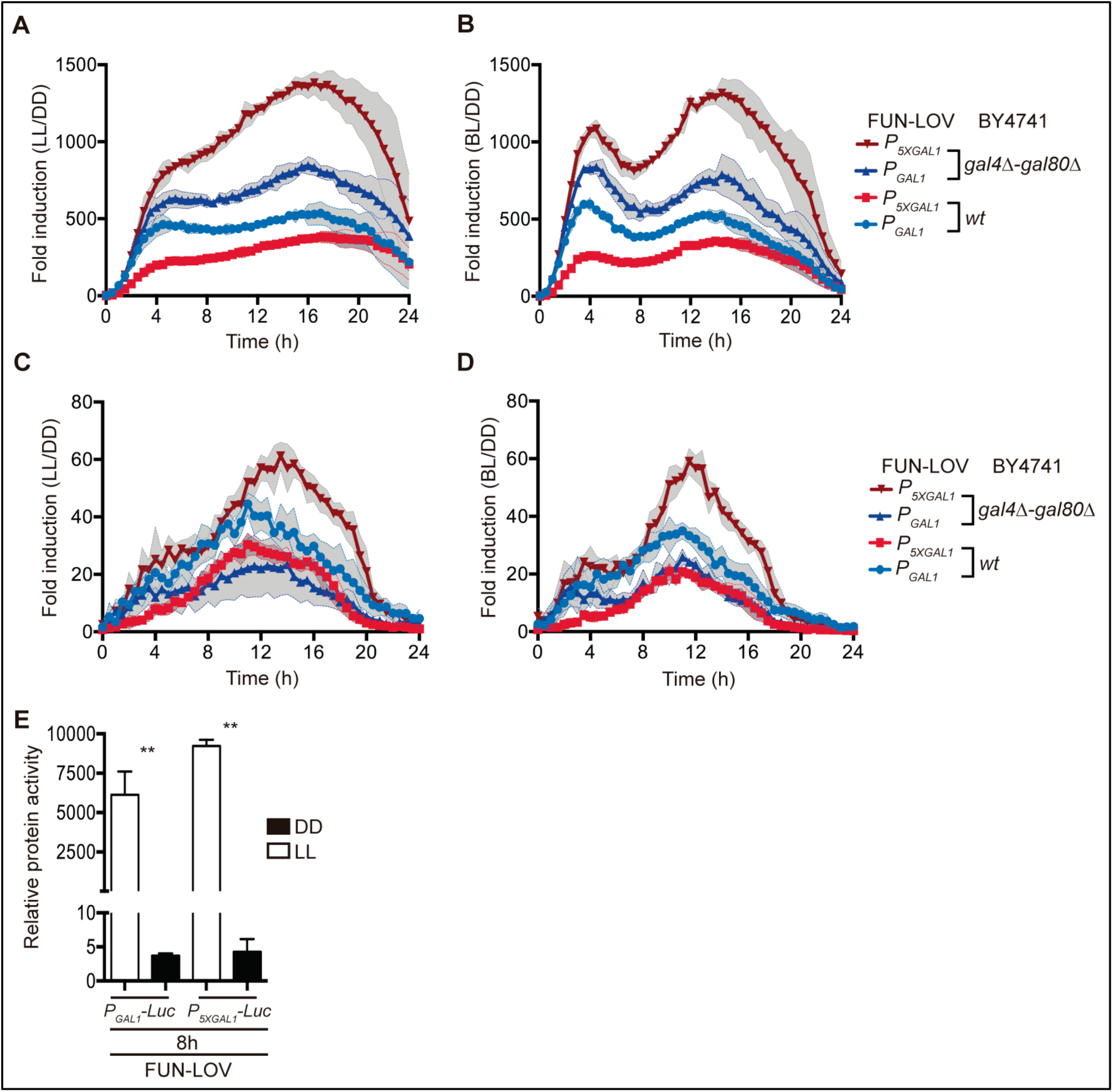
Levels of induction achieved by the FUN-LOV system. Fold of induction was calculated as the ratio of luciferase expression in constant white-light (LL) (A, C) or constant blue-light (BL) (B, D), respective to the average background expression in constant darkness (DD). As indicated (A, B), luciferase was episomally maintained under the control of two different promoters (*P*_*GAL1*_ and *P*_*5XGAL1*_) and was assayed in two different genetic backgrounds (BY4741 and BY4741 *gal4*Δ*-gal80*Δ). The ratio of luciferase activity was also calculated from strains bearing the luciferase inserted at the *GAL3* locus (C, D), under the control of *P*_*GAL1*_ and *P*_*5XGAL1*_ promoters, in two different genetic backgrounds, (BY4741 wt and BY4741 *gal4*Δ*-gal80*Δ). While in all experiments bioluminescence was measured in living cells, we also assayed luciferase activity from protein extracts, from cells harvested at 8 hours (8h) time points, observing approximately 1600 and 2500-fold of induction for *P*_*GAL1*_*-Luc* and *P*_*5XGAL1*_*-Luc*, respectively. The double asterisk (**) represents a significant statistical difference between LL and DD conditions (t-test, ** p<0.01) in panel (E). In panels (A) to (D), each plot corresponds to the average of six biological replicas with its standard deviation represented as shadowed regions. The luciferase activity assays were carried out in three biological replicas (n = 3), the average of these replicas with its standard deviation is shown in panel (E).

The feasibility to operate the FUN-LOV system as an on/off switch with dynamic and temporal resolution was also assayed. The FUN-LOV system yielded a distinct dynamic and temporal control of luciferase expression during the yeast exponential growth phase (Fig. 1F and Fig. S3). The system exhibited a decay in the stationary phase, probably resulting from the low transcriptional activity of the promoter (*P*_*ADH1*_) controlling the expression of the FUN-LOV components, which appeared as a common feature of the constitutive promoters evaluated in this work (Fig. S4). This also suggests that the expression levels reached by the FUN-LOV system might be further incremented by expressing its components under stronger promoters, such as *P*_*TEF1*_ or *P*_*TDH3*_ (Fig. S4C and S4D), which in the future could further boost the relative levels of the FUN-LOV switch components, such as the GAL4-DBD moiety (Fig. S4E). We also found that longer exposure to a blue-light pulse increased the expression of the reporter gene (Fig. 1G and Fig. S3E), which exemplifies the tuning of the response depending on the amount of provided light. However, an increase in the blue LED light intensity (from 20 to 40 μmol m^-2^ s^-1^) did not further augment luciferase levels (Fig. 1H and Fig. S3F), consistent with previous reports where intensities of 20 μmol m^-2^ s^-1^ or higher were considered high light fluency for WC-1/VVD interaction (26). Overall, the FUN-LOV system offers high expression levels with great dynamic and temporal resolution, making it a suitable switch for the control of biotechnologically phenotypes in yeast (27).

### FUN-LOV control of heterologous protein production

We then explored the capability of the FUN-LOV switch to regulate heterologous protein production upon light stimulation. Thus, the yeast codon optimized version of the *Cannabis sativa* Limonene Synthase (LS) gene was cloned in a pYES2 plasmid, allowing *P*_*GAL1*_ promoter control and V5 tagging of the protein for Western blot analysis (Fig. 3A). This protein is involved in the production of limonene, a compound widely used by the food and house hold industries to increase lemon scent in their products (28). Western blot analysis showed that yeast cells expressing the FUN-LOV system achieved a LS induction of 110-fold (LL/DD), which is 2.5 times higher than the average induction (44-fold) achieved, in our hands, by galactose. Notably, FUN-LOV yields lower background levels in DD conditions, particularly compared to the LS levels in the OFF state of the galactose/glucose systems (Fig. 3B, 3C and 3D). Furthermore, when we evaluated the effect of LL on the temporal expression of proteins by exposing the cells to 2-hour pulses of white-light or blue-light, high levels of protein expression were obtained, with no differences between both light sources in two different genetic backgrounds (Fig. S5). Overall, FUN-LOV allows high levels of heterologous protein expression upon light stimulation, with low background expression in darkness, and reaching induction levels surpassing chemical induction approaches.

**FIG 3.**
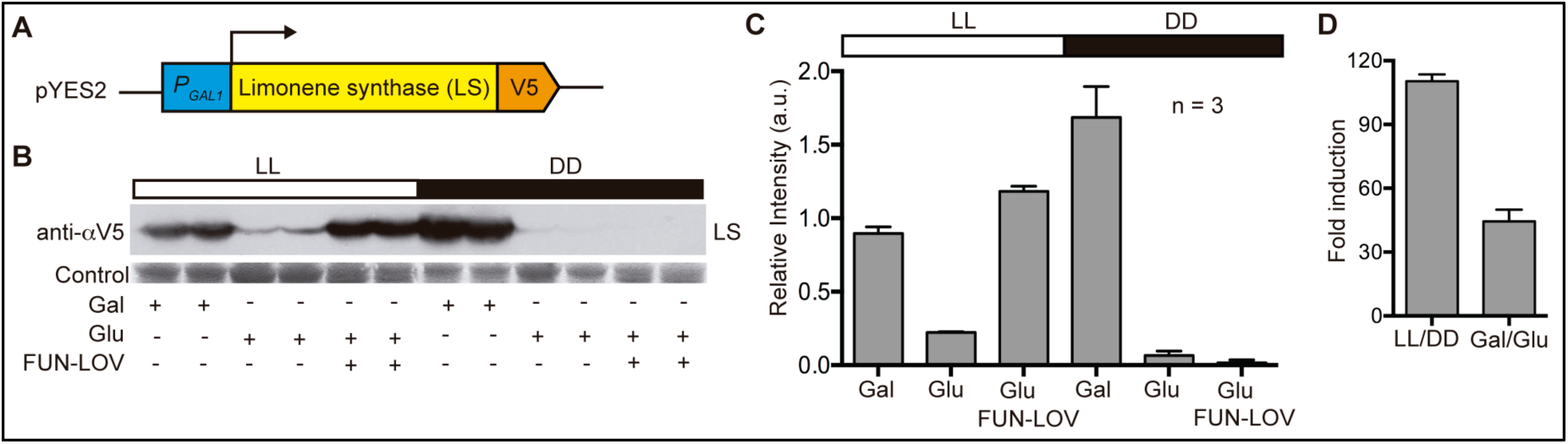
Heterologous protein expression controlled by the FUN-LOV switch. (A) The cDNA encoding for limonene synthase (LS) was used for its synthesis with codon optimization for yeast and cloning in a pYES2 expression plasmid, allowing LS control by *P*_*GAL1*_ promoter and including a C-terminal V5-tag for Western blot detection. (B-C) LS expression was analysed by Western blot (B) and normalized to membrane stained proteins (C), under constant white-light (LL) and constant darkness (DD) conditions. Strains carrying the pYES-LS plasmid reveal strong expression of LS by Galactose (Gal), but not glucose (Glu), as expected from the control of the endogenous Gal4 TF over a Gal1 promoter. If, in addition to the pYES2-LS plasmid the cells include the FUN-LOV switch, expression is tightly regulated by light. (D) Comparison of Gal/Glu and LL/DD fold of inductions, the FUN-LOV systems reveals higher fold of expression relative to a classic Gal expression protocol. The Western blot experiments were performed in three biological replicas (n = 3), the average of these replicas with its standard deviation is shown in panels (C) and (D).

### Modulation of yeast flocculation by light

Finally, we sought to apply the FUN-LOV system to optogenetically modulate a biotechnologically relevant process such as flocculation. We rationalized that direct overexpression of the main genes responsible for flocculation (*FLO1* or *FLO11*) would trigger such phenotype, in agreement with previous reports for these genes (23, 29). On the other hand, low expression of the yeast transcriptional co-repressor *TUP1* should lead to upregulation of its targets (30), among which is *FLO1*, therefore triggering strong flocculation. Initially, we swapped the endogenous promoters of *FLO1, FLO11* and *TUP1* genes by *P*_*GAL1*_ and *P*_*5xGAL1*_ promoters (Fig. 4A). We confirmed the correct replacement of the endogenous promoters and analysed the behaviour of the resulting strains by growing them in galactose and glucose conditions, as carbon sources. Using bright-field microscopy, we observed strong cell aggregation in strains carrying *P*_*GAL1*_*-FLO1* in galactose but not in glucose; but moderate aggregation in the ones bearing *P*_*GAL1*_*-FLO11* (Fig. S6). As expected, strains carrying *P*_*GAL1*_*-TUP1* showed strong cell aggregation in glucose but not in galactose, confirming the flocculation phenotype caused by the lack of *TUP1* expression (Fig. S6). Then, we episomally incorporated the FUN-LOV system into the strains carrying the swapped promoters. The strains with *P*_*GAL1*_-*FLO1* and carrying the FUN-LOV system demonstrated high levels of cell aggregation in LL; and low or no aggregation in DD (Fig. S7). We denominated this FUN-LOV-controlled phenotype as Flocculation in Light (FIL). Compared to the FUN-LOV control of *FLO1* driven flocculation, only a modest FIL phenotype was observed in LL for strains with *P*_*GAL1*_*-FLO11.* These phenotypes were confirmed by macroscopic observation, as well as by fluorescence microscopy of yeast cells constitutively expressing *mCherry*, and by calculating the flocculation index of each strain (Fig. 4B and 4C). On the other hand, strains with FUN-LOV control of *P*_*GAL1*_*-TUP1* revealed high levels of cell aggregation in DD, but not in LL (Fig. 4B, 4C and Fig. S7). This phenotype was denominated Flocculation in Darkness (FID).

**FIG 4.**
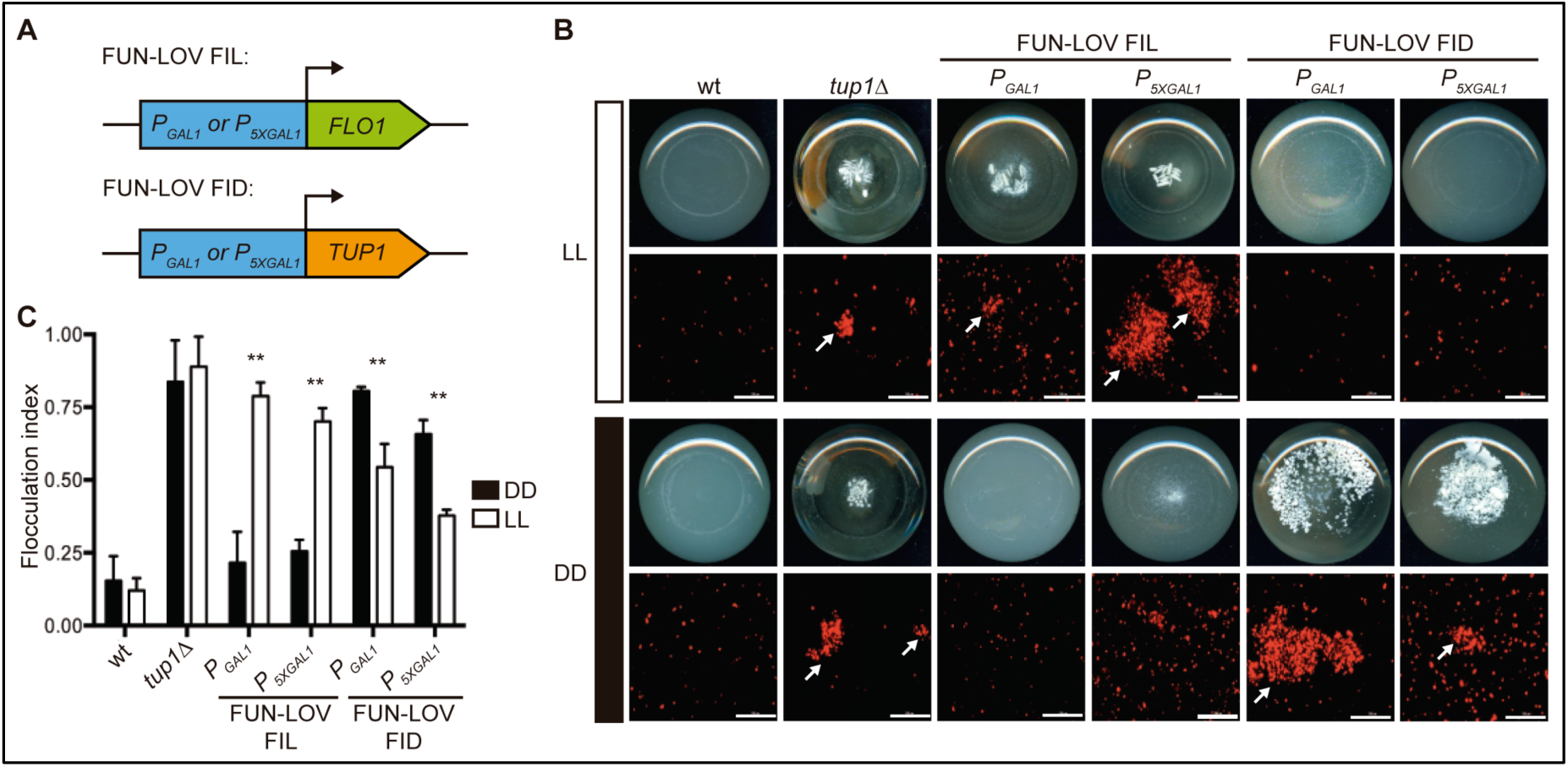
The FUN-LOV switch allows precise control of Flocculation by light conditions. (A) FUN-LOV dependent expression of *FLO1* or *TUP1* genes allows flocculation to occur in Light (FIL) or in darkness (FID). (B) FIL and FID phenotypes observed as macroscopic cell aggregation in culture flasks (bottom view) and fluorescence microscopy of cells expressing *mCherry* under the *P*_*TDH3*_ constitutive promoter. Experiments were performed under constant white-light (LL) or constant darkness (DD) conditions. The BY4741 wild type (wt) and BY4741 *tup1*Δ strains were utilized as negative and positive controls of flocculation, respectively. Cell aggregates are exemplified with arrows, scale bar represent 100 µm with a 20X objective using a Cytation 3 in fluorescence microscope mode. (C) Quantification of the FIL and FID phenotypes observed in panel (B) by calculating the flocculation index, computed as the OD difference recorded in the yeast culture after 30 min of static incubation: 1-(Final_OD600_/Initial_OD600_). Statistical significant differences between LL and DD conditions are indicated as ** (t-test, p<0.01). The flocculation experiments were conducted in three biological replicas (n = 3), the average of these replicas with its standard deviation is shown in panel (C).

The reversibility of the flocculation phenotype was evaluated under different conditions of LL and DD. Growth of *FLO1* FIL strains during 24 hours in DD followed by 24 hours in LL resulted in strong flocculation (Fig. 5A and 5B), whereas the opposite treatment - 24 hours in LL followed by 24 hours in DD – revealed strong flocculation in LL without reversion of the phenotype after the transfer to DD (Fig. 5C and 5D). In the case of *TUP1* FID strains, there was no flocculation in LL during the initial 24 hours of incubation, followed by strong flocculation after the transfer to DD (Fig. 5E and 5F). As expected, no reversion was observed for the phenotype when cells were incubated 24 hours in DD and then transferred to LL (Fig. 5G and 5H). Finally, we evaluated the time necessary to induce the FIL and FID phenotypes in yeast cultures. The results showed that the FIL phenotype becomes readily visible after 6 hours of light stimulation, whereas the FID phenotype can be observed after 10 hours of incubation in the dark (Fig.5I). Overall, depending on the genetic wiring of FUN-LOV, yeast flocculation can be triggered by light (FIL phenotype) or by its absence (FID phenotype). These phenotypes showed no reversibility and the time of its appearance had different temporalities.

**FIG 5.**
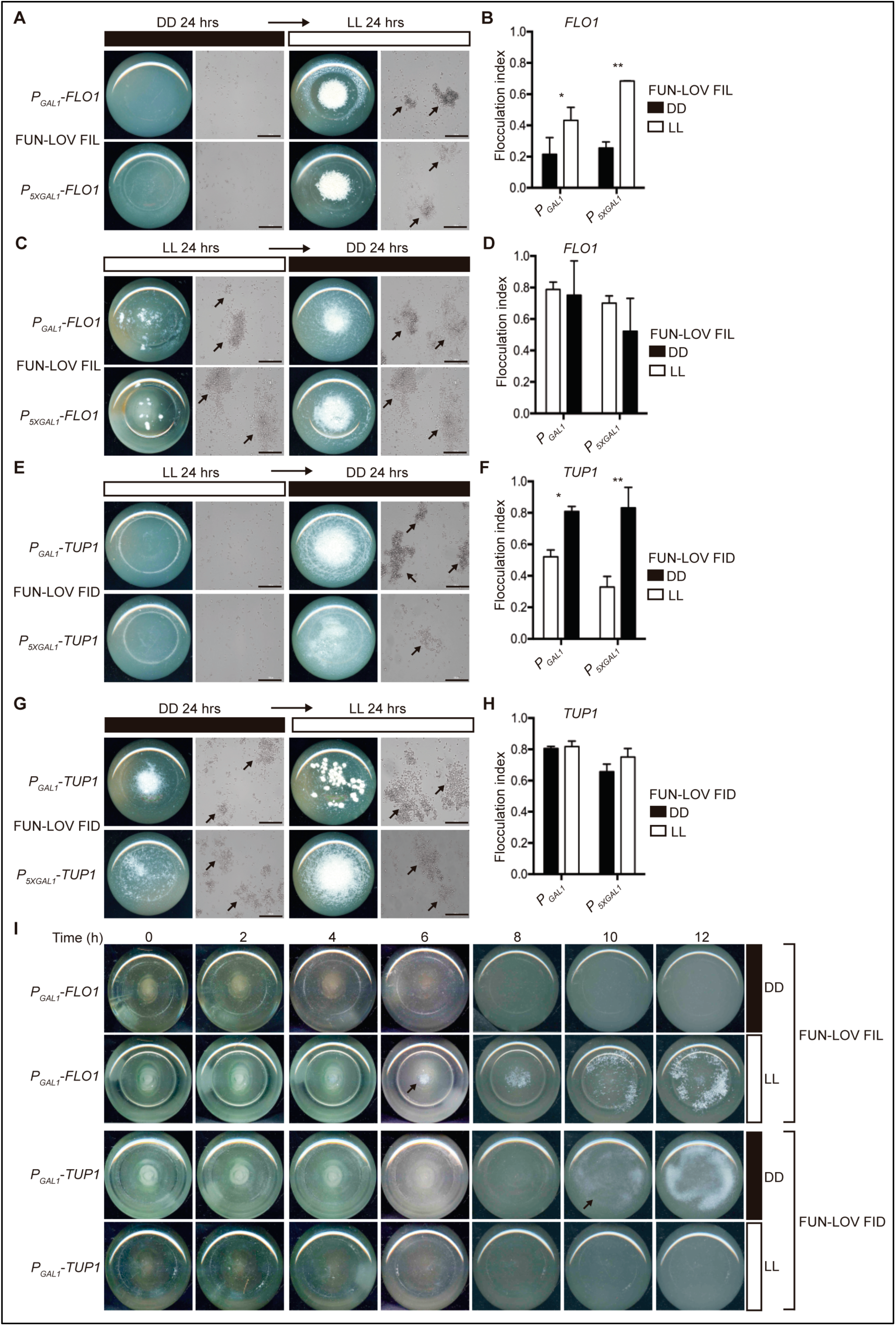
Reversibility and temporality of the different flocculation phenotypes. (A-D) The reversion of the FIL phenotype for strains with *FLO1* promoter swapping and carrying the FUN-LOV system, was assayed after 24 hours of incubation in constant darkness (DD) and then transferred 24 hours to constant light (LL) (A-B); the opposite regimen was also assayed with 24 hours in LL and 24 hours in DD (C-D). Cellular aggregation was recorded with pictures of the bottom view of the culture flasks and bright-field microscopy (A and C). Yeast aggregates are highlighted with arrows, scale bar represent 100 µm with a 20X objective using a Cytation 3 in microscope mode. The flocculation index was also assayed for the same set of strains and culture conditions (B and D). Single (*) and double asterisk (**) represents statistically significant differences between LL and DD conditions (t-test, * p<0.05 and ** p<0.01). (E-H) Reversion of the FID phenotype for strains with *TUP1* promoter swapping and carrying the FUN-LOV system, was also assayed under the same experimental conditions of panels (A) to (D), starting from non-flocculating condition (LL) to the flocculating condition (DD) (E); the opposite regimen was also assayed (H). (I) Time course of the FIL and FID phenotypes for the strains with *FLO1* and *TUP1* promoter swapping (by *P*_*GAL1*_) and carrying the FUN-LOV system. The cellular aggregation was recorded with pictures of the bottom view of the culture flasks every two hours in DD and LL conditions. The arrows highlight the time point where the macroscopic flocculation phenotype started to be visible. The flocculation experiments were performed in three biological replicas (n = 3), the average of these replicas with its standard deviation is shown in panels (B), (D), (F) and (H).

## DISCUSSION

Based on the LOV-domains of two blue-light photoreceptors from the fungus *N. crassa* (proteins WC-1 and VVD), we developed a new optogenetic switch named FUN-LOV. We set-up FUN-LOV in yeast cells, since this microorganism is a model eukaryotic system for genetics and molecular biology studies, and a powerful platform for biotechnology, including applications such as production of high value metabolites and proteins (31, 32). Using yeast as biological chassis, FUN-LOV showed three singular features: 1, high levels of gene expression with a broad dynamic range and temporal resolution (Fig. 1D, 1E and 1F); 2, low background expression in dark conditions (Fig. 1C and Fig. 2); 3, the control of biotechnologically relevant phenotypes in yeast, such as heterologous protein expression (Fig. 3) and flocculation (Fig. 4 and 5). Those characteristics achieved by FUN-LOV increase its potential uses in industrial or high-volume bioprocess conditions.

The first optogenetic switch developed and implemented in yeast was based on the red-light dependent interactions of the photoreceptor Phytochrome B (PhyB) and its interacting protein PIF3 from *Arabidopsis thaliana* (6). Ever since, this switch has been adapted in yeast with different purposes such as: reconstitution of protein activities (7), subcellular protein localization, control of cell polarity and budding phenotypes (33, 34), and reporter gene expression (YFP or GFP) (19, 35). Recently, a new red-light optogenetic switch also based on PhyB-PIF3 interaction (named PhiReX), that overcomes the obstacle of external addition of the chromophore phycocyanobilin (PCB), has been implemented and tested in yeast (35). These types of toggle switches can be quite powerful as they provide activation with a given wavelength (red) to then be turned off by another one (far-red). Yet, in some cases that contrasts with the simplicity of blue-light switches which can be readily activated by blue or white light, while turned off with fast kinetics upon transfer to the dark, without the need extra specific wavelengths.

In general, multiples blue-light optogenetic switches have been implemented in several biological platforms utilizing a repertoire of biological parts, resulting in diverse expression levels. The latter has been monitored by different reporter genes, a fact that complicates the direct comparison between systems (22). In yeast, one of the most widely used switch is based on the blue-light dependent interaction of the photoreceptor Cryptochrome 2 (CRY2) and its interacting protein CIB1 from *A. thaliana*, allowing light-controlled reporter gene expression (20), heterologous expression of YFP or mCherry (36, 37) and cell cycle modulation (19). Besides cryptochromes, blue-light photoreceptors containing LOVs domains have been also used in yeast optogenetics. The LOV2 domain from *Avena sativa* phototropin 1 (AsLOV2) was utilized for light-controlled caging of peptides and ß-galactosidase expression in yeast (38). Similarly, the LOV2 domain from *A. thaliana* phototropin 1 (AtLOV2) has been also used for blue-light control of the cell cycle progression and expression of conditional essential genes (39, 40).

Fungal blue-light photoreceptors containing LOV domains have been seldom utilized in optogenetics and its uses have not been exploited in yeast, regardless the functional conservation between fungal and plant LOV-domains (41). Solely the protein VIVID (VVD) from the fungus ***N***. *crassa* has been used in optogenetics switches, permitting light-controlled expression of transgenes in mice and mammalian cells (13, 14), subcellular protein localization (42) and genome engineering in cell lines (43). In ***N***. *crassa*, the protein VVD not only has the capacity to self-dimerize upon light stimulation, but additionally VVD interacts with the protein WC-1, a LOV-LOV interaction that allows photoadaptation to different light intensities (17, 44). Indeed, WC-1 is a GATA transcription factor and also a blue-light photoreceptor containing a LOV domain, participating (with WC-2 protein) in the White-Collar Complex (WCC), which acts as positive element in the circadian clock of ***N***. *crassa* (45). Therefore, when it comes to LOV, finding the right partner may improve the performance of the system, as evidence by the VVD/WC-1 combination. Considering the diversity of such sequences in different fungal genomes, such LOV domains are a promising source for novel optogenetic switches as the one presented herein.

In conclusion, in this work we reported the implementation in yeast of a novel optogenetic switch, FUN-LOV, which provides accurate and strong light-controlled gene expression, exemplified also by the regulation of two phenotypes of biotechnological relevance, such as heterologous protein expression and flocculation. The current challenge is to successfully scale up of this technology for industrial bioprocesses, under conditions where high culture densities could preclude efficient light delivery to all yeast cells. Importantly, in addition to its applicability to produce high value metabolites or heterologous proteins, its low background and broad dynamic range makes FUN-LOV a powerful tool to exquisitely regulate the expression of any gene of interest and to probe complex biological phenomena. In addition, its modular design can help the implementation of different opto-logic gates, by combining it with optogenetic switches for other light wavelengths (22), in order to rewire, built or perturb complex gene regulatory networks in yeast. Notably, as FUN-LOV utilizes a Gal4-DBD, it could be readily domesticated in other systems such as mammals, or Drosophila, where Gal4-orthogonal control of gene expression has proven extremely successful.

## MATERIALS AND METHODS

### Yeast strains, media and culture conditions

*Saccharomyces cerevisiae* strains BY4741 *wt* (*MATa; his3*Δ*1; leu2*Δ*0; met15*Δ*0; ura3*Δ*0*) and BY4741 with *GAL4* and *GAL80* deletions (*MATa; his3*Δ*1; leu2*Δ*0; met15*Δ*0; ura3*Δ*0, gal4*Δ*::NatMx, gal80*Δ*::HphMx*), were used as genetic background for gene deletions and promoter swapping. The strains used and generated in this work were maintained in YDPA medium (2% glucose, 2% peptone, 1% yeast extract and 2% agar) at 30 °C and their genotypes are listed in Table 1. Strains carrying plasmids with auxotrophic markers were maintained in synthetic complete (Sc) media (0.67% yeast nitrogen base without amino acids, glucose 2%, dropout mix 0.2% and agar 2%) minus the corresponding amino acid (dropout mix). Growth of yeast cultures under different white-light (LL) and darkness (DD) conditions were conducted in Percival incubators (Percival Scientific, USA) at 30 °C, using 100 μmol m^2^ s^-1^ of white-light intensity. The blue-light (BL) experiments were carried out using a LED lamp with 20 μmol m^2^ s^-1^ and 40 μmol m^2^ s^-1^ of light intensity. The specific wavelength of the blue light is 460 nm. In general, cells were grown in 50 mL of Sc media at 30 °C with 130 RPM of shaking in flasks or in 200 µL of Sc media using 96-well plates at 30 °C. Manipulation of strains in the “dark” was conducted in a light-tight room equipped with red safety lights.

**Table 1.** Strains of *Saccharomyces cerevisiae* used and generated in this work.

| Strain | Genotype | Source |
| --- | --- | --- |
| BY4741 | MA Ta his3Δ1 leu2Δ0 met15Δ0 ura3Δ0 | Euroscarf |
| gal4Δ/gal80Δ | BY4741; gal4Δ::NatMx gal80Δ::HphMx | This work |
| flo1Δ | BY4741; flo1Δ::URA3 | This work |
| flo11Δ | BY4741; flo11Δ::URA3 | This work |
| tup1Δ | BY4741; tup1Δ::KanMx | Euroscarf |
| P<sub>GAL1</sub>-FLO1 | BY4741; P<sub>FLO1Δ</sub>::KanMxRV-P<sub>GAL1</sub> | This work |
| P<sub>5XGAL1</sub>-FLO1 | BY4741; P<sub>FLO1Δ</sub>::KanMxRV-P<sub>5XGAL1</sub> | This work |
| P<sub>GAL1</sub>-FLO11 | BY4741; P<sub>FLO11Δ</sub>::KanMxRV-P<sub>GAL1</sub> | This work |
| P<sub>5XGAL1</sub>-FLO11 | BY4741; P<sub>FLO11Δ</sub>::KanMxRV-P<sub>GAL1</sub> | This work |
| P<sub>GAL1</sub>-TUP1 | BY4741; P<sub>TUP1Δ</sub>::KanMxRV-P<sub>GAL1</sub> | This work |
| P<sub>5XGAL1</sub>-TUP1 | BY4741; P<sub>TUP1Δ</sub>::KanMxRV-P<sub>5XGAL1</sub> | This work |
| gal3Δ::P<sub>GAL1</sub>-Luc | BY4741; gal3Δ::KanMxRV-P<sub>GAL1</sub>-Luc | This work |
| gal3Δ::P<sub>5XGAL1</sub>-Luc | BY4741; gal3Δ::KanMxRV-P<sub>5XGAL1</sub>-Luc | This work |
| gal4Δ/gal80Δ<br>P<sub>GAL1</sub>-FLO1 | BY4741; gal4Δ/gal80Δ; P<sub>FLO1Δ</sub>::KanMxRV-P<sub>GAL1</sub> | This work |
| gal4Δ/gal80Δ<br>P<sub>GAL1</sub>-FLO11 | BY4741; gal4Δ/gal80Δ; P<sub>FLO11Δ</sub>::KanMxRV-P<sub>GAL1</sub> | This work |
| gal4Δ/gal80Δ<br>P<sub>GAL1</sub>-TUP1 | BY4741; gal4Δ/gal80Δ; P<sub>TUP1Δ</sub>::KanMxRV-P<sub>GAL1</sub> | This work |
| gal4Δ/gal80Δ<br>gal3Δ::P<sub>GAL1</sub>-Luc | BY4741; gal4Δ/gal80Δ; gal3Δ::KanMxRV-P<sub>GAL1</sub>-Luc | This work |
| gal4Δ/gal80Δ<br>gal3Δ::P<sub>5XGAL1</sub>-Luc | BY4741; gal4Δ/gal80Δ; gal3Δ::KanMxRV-P<sub>5XGAL1</sub>-Luc | This work |

### Plasmid constructs and strains generation

Plasmids carrying the LOV domains of WC-1 and VVD photoreceptors were generously provided by the Brunner Lab (17). The plasmids used and generated in this work are shown in the Table 2. The FUN-LOV components were cloned into pRS423 (GAL4-DBD plus WC-1) and pRS425 (GAL4-AD plus VVD) plasmids for *HIS3* and *LEU2* auxotrophic selection, respectively. All the cloning experiments and genetic constructs were generated using yeast recombinational cloning in-vivo assembly (46, 47).

**Table 2.** Plasmids assembled in this work using yeast recombinational cloning.

| Plasmid | Construct |
| --- | --- |
| pRS423-WC1 | *P<sub>ADH1</sub>-WC1 LOV-GAL4 DBD-ADH2<sub>ter</sub>* |
| pRS425-VVD | *P<sub>ADH1</sub>-VVD36-GAL4 AD-ADH2<sub>ter</sub>* |
| pRS426-*P<sub>GAL1</sub>-Luc* | *KanMxRV-P<sub>GAL1</sub>-Luc-CYC1<sub>ter</sub>* |
| pRS426-*P<sub>5XGAL1</sub>-Luc* | *KanMxRV-P<sub>5XGAL1</sub>-Luc-CYC1<sub>ter</sub>* |
| pRS426-*P<sub>GAL1</sub>* | *KanMxRV-P<sub>GAL1</sub>* |
| pRS426-*P<sub>5XGAL1</sub>* | *KanMxRV-P<sub>5XGAL1</sub>* |
| pRS426-*mCherry* | *P<sub>TDH3</sub>-mCherry-CYC1<sub>ter</sub>-HphMx* |
| pRS426-*P<sub>ADH1</sub>-Luc* | *P<sub>ADH1</sub>-Luc-CYC1<sub>ter</sub>* |
| pRS426-*P<sub>TEF1</sub> Luc* | *P<sub>TEF1</sub>-Luc-CYC1<sub>ter</sub>* |
| pRS426-*P<sub>TDH3</sub>-Luc* | *P<sub>TDH3</sub>-Luc-CYC1<sub>ter</sub>* |

The synthetic version of the *P*_*GAL1*_ promoter, called *P*_*5xGAL1*_, carrying five DNA binding sites for the Gal4 transcription factor (GAL4-UAS) was synthetized using Genewiz gene synthesis service. In the constructs assembly, the *P*_*GAL1*_ and *P*_*5xGAL1*_ promoters were amplified by PCR using Phusion Flash High-Fidelity PCR Master Mix (Thermo scientific, USA). Additionally, the Kanamycin (*KanMx*) antibiotic resistance cassette was added in the reverse direction upstream of each promoter (*KanMxRv-P*_*GAL1*_ or *KanMxRv-P*_*5xGAL1*_) using yeast recombinational cloning (47). The complete genetic construct (*KanMxRv-P*_*GAL1*_ or *KanMxRv-P*_*5xGAL1*_) was used to transform the BY4741 *wt* and BY4741 *gal4*Δ*-gal80*Δ strains. The complete genetic constructs (*KanMxRv-P*_*GAL1*_ or *KanMxRv-P*_*5xGAL1*_) were amplified by PCR using a Phusion Flash High-Fidelity PCR Master Mix (Thermo scientific, USA) and 70 bp primers for direct homologous recombination on the target locus, allowing the swapping of the endogenous promoter region. The promoter swapping of *FLO1, FLO11* and *TUP1* were confirmed by PCR under standard conditions, primers used for plasmids assembly, promoter swapping and promoter swapping confirmations are shown in Table S1. The same assembly procedure was followed to construct different versions of the luciferase reporter gene under the control of *P*_*GAL1*_ and *P*_*5xGAL1*_ promoters. The deletion of *GAL4, GAL80* and luciferase reporter gene integrations *GAL3* locus were carried out using one-step PCR deletion by recombination (48). Primers used for gene deletion and reporter gene integration in the genome and its confirmations are listed in Table S1.

### Luciferase in vivo expression and Western blot assay

We used a previously described destabilized version of the firefly luciferase for real-time monitoring of gene expression in yeast (49). We cloned into plasmid pRS426 the luciferase reporter gene under the control of *P*_*GAL1*_ and *P*_*5xGAL1*_ promoters using yeast recombinational cloning as we described above. Real-time luciferase expression was measured under DD, LL and BL conditions using a Cytation 3 microplate reader (BioTek, USA), which allows the measurement of both, OD_600_ and luminescence of the cell cultures over time. Briefly, the yeast strains were grown overnight in a 96 well plate with 200 uL of Sc medium at 30 °C in DD condition, 10 uL of these cultures were used to inoculate a new 96 well plate containing 290 uL (30-fold dilution) of fresh media plus 1 mM of luciferin. The OD_600_ and the luminescence were acquired every 30 minutes using a Cytation 3 microplate reader, running a discontinuous kinetic protocol with 30 seconds of shaking (285 cycles per minute) before each measurement. This protocol also allowed illumination of the 96 well plate between each measurement, keeping the plate outside of the equipment and exposing it to the light source. Luciferase expression was normalized by OD (600 nm) of the yeast cultures and all experiments were performed in six biological replicas (50). Normalization of real-time luciferase levels was necessary as yeast biomass was rapidly changing over the length of the experiment, something that may not be necessary when monitoring luciferase from filamentous fungi (51).

Limonene synthase enzyme cDNA sequence from *Cannabis sativa* was codon optimized for *S. cerevisiae* and cloned into pYES2 expression vector. The strain BY4741 *wt* was co-transformed with pYES2-LS plasmid and the components of the FUN-LOV system. Strains were grown in DD and LL conditions until OD_600_ = 1, protein extractions of the yeast cellular pellets were carried out standard conditions (52). We used 25 ug of total protein from each sample for western blot assays, which were conducted in three biological replicas with a α-V5 antibody (Invitrogen, USA), using a 1/5000 dilution (51, 53).

### Gene expression by real-time PCR (qPCR)

The gene expression levels for the FUN-LOV component (GAL4-DBD) was measured by qPCR in the BY4741 genetic background under LL condition. Briefly, the yeast strains were grown during 8 hours in LL condition at 30 °C in Sc media for RNA extractions. RNA extractions were carried out using Tryzol Reagent (Thermo Fisher Scientific, USA) and 100 ng of total RNA were used for reverse transcription using Superscript III transcriptase (Invitrogen, USA) under standard conditions. The cDNA was amplified using 2X SensiMix SYBR Hi-ROX Kit (Bioline, USA) and using a StepOnePlus Real-Time PCR equipment (Thermo Fisher Scientific, USA). The relative gene expression was calculated using the 2^-ΔΔCt^ method and utilizing two different reference genes (*HEM2* and *TAF10*) (54, 55). The primers used for qPCR are listed in Table S1.

### Luciferase enzymatic activity

The luciferase protein activity was assayed using the Luciferase Assay System kit (Catalog number E1500, Promega, USA) with modifications. Briefly, the yeast strains were grown during 8 hours at 30 °C in Sc media and harvest by centrifugation during 5 min at 4000g. The cell pellet was disrupted using 200 uL of 2x Lysis Buffer Reagent (Promega, USA) and 200 uL of glass beads in a TissueLyser II equipment (QIAGEN, USA) during 3 minutes. Cells were centrifuged 5 min at max speed and the supernatant containing the protein extract was recovered and quantified by the Bradford standard method. The luciferase activity was assayed combining 5 µL of the total extracted proteins plus 100 µL of Luciferase Assay Reagent (Promega, USA). The luminescence was immediately recorded in a Cytation 3 microplate reader (BioTek, USA) and it was normalized using the total protein concentration of each sample.

### Flocculation phenotypes

Strains with promoter swapping in *FLO1, FLO11* and *TUP1* and carrying the FUN-LOV system were evaluated under DD and LL conditions as we previously described. Scanning of the flocculation phenotype were taken after 24 hours of growth in culture flasks under DD or LL conditions. Additionally, the strains were transformed with a pRS426 plasmid carrying *mCherry* controlled by the *P*_*TDH3*_ promoter. Pictures of yeast cells were taken under bright-field and fluorescence microscopy using a Cytation 3 in microscope mode (BioTek, USA). In the time course experiments scanning of the culture flask was performed every two hours. The flocculation of each strain was quantified calculation the flocculation index of each strain at OD 600 nm (56, 57). The flocculation index was calculated as the OD difference in a yeast culture after 30 min of static incubation: 1-(Final_OD_/Initial_OD_).

## ACKNOWLEDGEMENTS

We thank Consuelo Olivares, Rodrigo Santibañez and Cristobal Mena for technical help during this work, and members of the Larrondo’s lab for their constructive comments on this manuscript. This research was funded by MIISSB Iniciativa Científica Milenio-MINECON, CONICYT/FONDEQUIP EQM130158, and CONICYT/FONDECYT 1171151 & 1170745 to L.F.L. and E.A. respectively. F.S. was supported by CONICYT/FONDECYT (grants no. 3150156 and 11170158), V.R and V.D were supported by CONICYT/PhD scholarships 21170331 and 6313018 respectively. L.F.L is an International Research Scholar of the Howard Hughes Medical Institute.

## AUTHOR CONTRIBUTIONS

F.S., V.R., V.D., E. A. and L.F.L. designed research; F.S., V.R., V.D. and J.L. performed lab experiments and analysed the experimental data; F.S. and L.F.L. wrote the paper with insight from all the authors.

