## Supplementary Materials for "FUN-LOV: Fungal LOV domains for optogenetic control of gene expression and flocculation in yeast"

### Supporting Figures

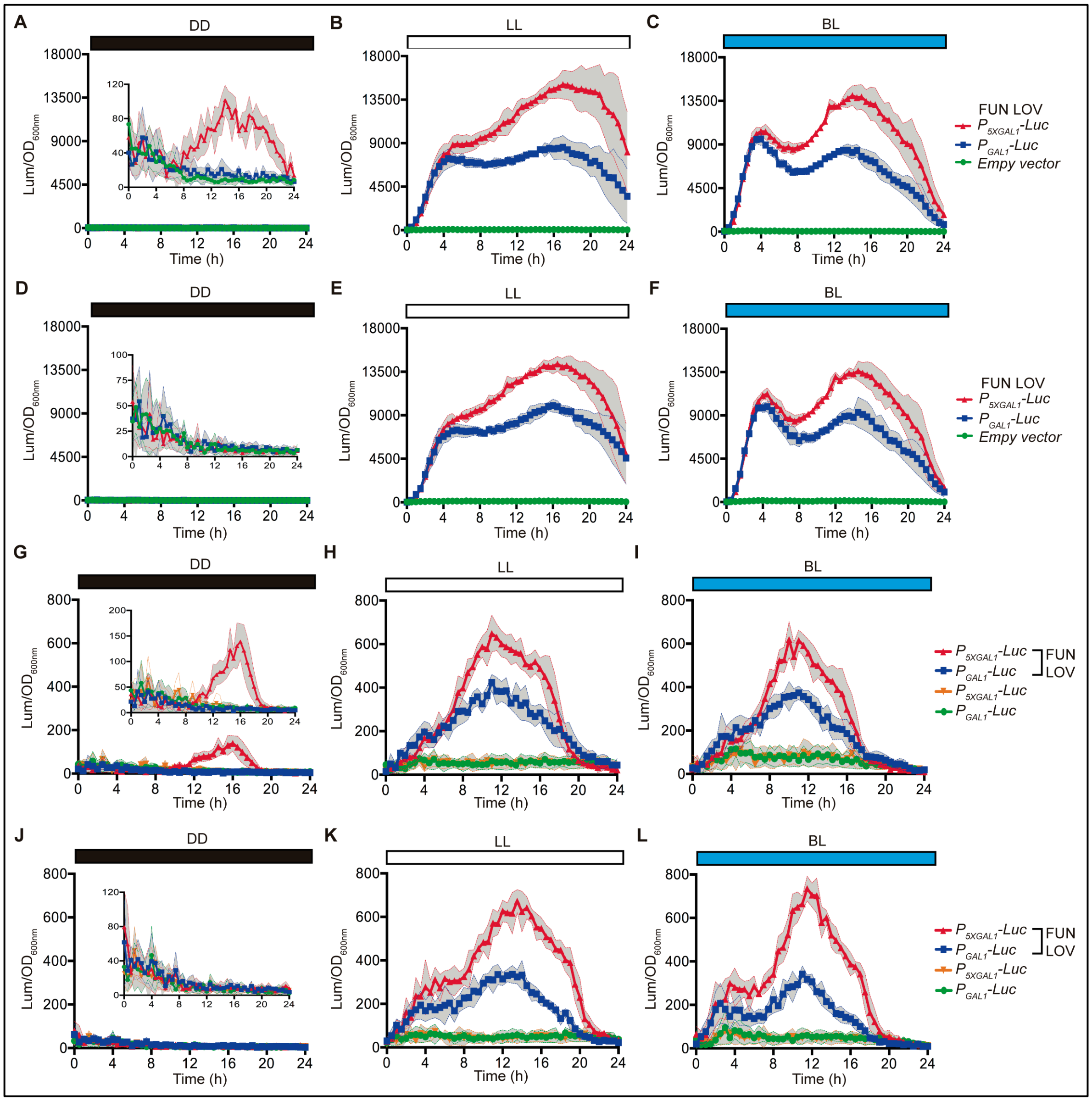


**FIG S1. FUN-LOV control of luciferase expression.** Bioluminescence was monitored and normalized by $OD_{600}$, as cultures were grown in constant darkness (DD), constant white-light (LL) or constant blue-light (BL), respectively. Components of the FUN-LOV system and the reporter construct ($P_{GAL1}$-luciferase or $P_{5XGAL1}$-luciferase) were maintained episomally in a BY4741 wild type (A-C), or in a BY4741 *gal4Δ-gal80Δ* strain (D-F), as the latter helps reducing background expression in DD. BY4741 (G-I), or BY4741 *gal4Δ-gal80Δ* -derived strains (J-L), containing the FUN-LOV system episomally, but having the luciferase reporters integrated at the *GAL3* locus were also evaluated. In panels (A) to (L)**,** each plot corresponds to the average of six biological replicas with its standard deviation represented as shadowed regions.

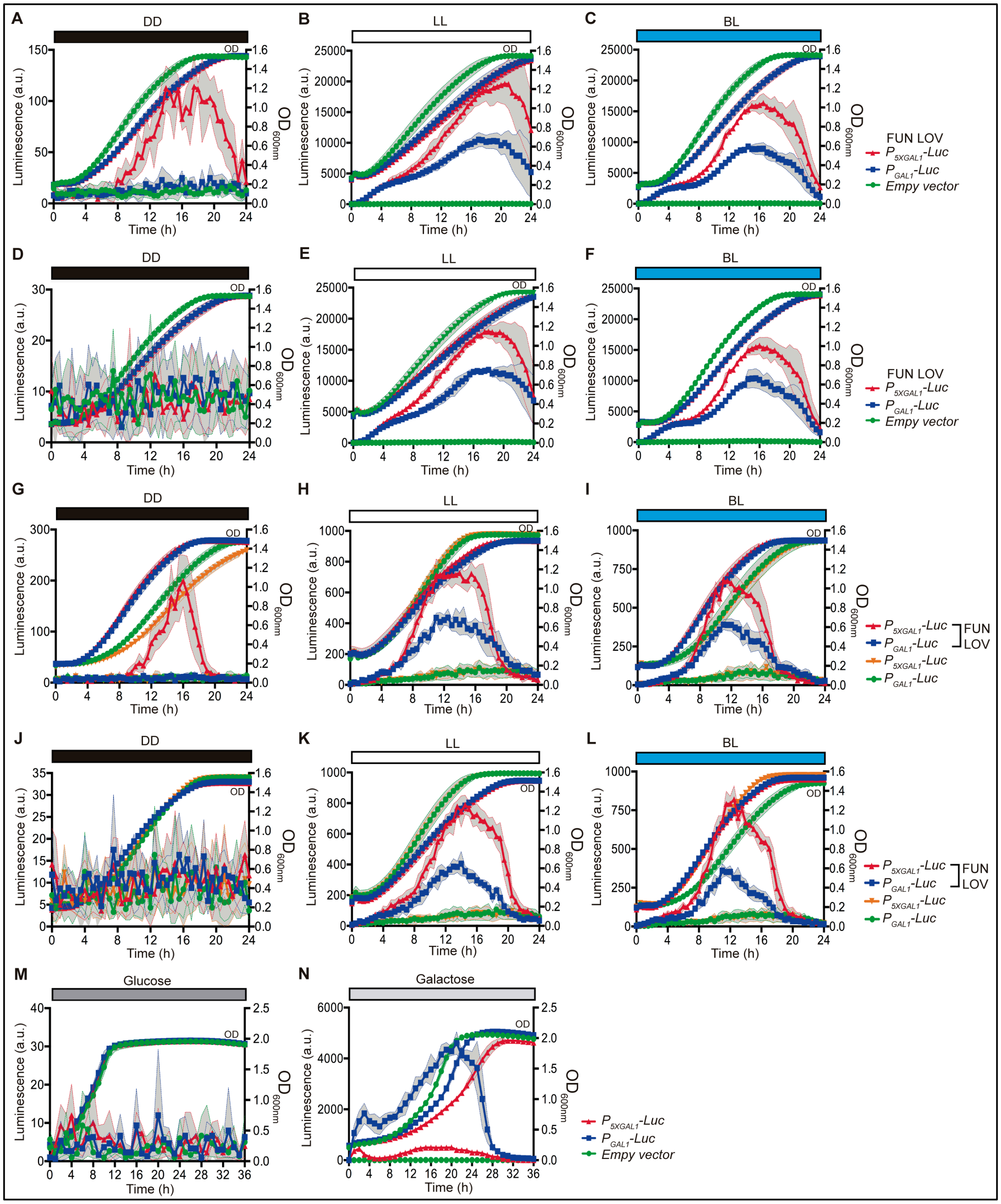


**FIG S2. Raw luciferase expression levels during active yeast growth.** (A-C) The growth of yeast strains containing FUN-LOV and luciferase controlled by a $P_{GAL1}$ or $P_{5XGAL1}$ promoter was monitored as $OD_{600nm}$ and bioluminescence levels, which were quantified in constant darkness (DD), constant white-light (LL) or constant blue-light (BL), respectively. Components of the FUN-LOV switch and the reporter gene were maintained episomally in BY4741 wild type (A-C) or BY4741 *gal4*Δ-*gal80*Δ strain (D-F). (G-L) Although the FUN-LOV switch was kept episomally, the luciferase reporter was chromosomally inserted at the *GAL3* locus in BY4741 (G-I) or in the

BY4741 *gal4*Δ-*gal80*Δ (J-L) background. In addition, the behaviour of the episomal Luc-reporter (in a BY4741 background, and depending on the endogenous Gal4p, was evaluated in glucose (M) or galactose (N), which resembles DD (no-expression) and LL (inducing conditions). When indicated, raw data for luciferase expression was acquired from two different type of promoters: $P_{GAL1}$ and $P_{5XGAL1}$. In panels (A) to (N), each plot corresponds to the average of six biological replicas with its standard deviation represented as shadowed regions.

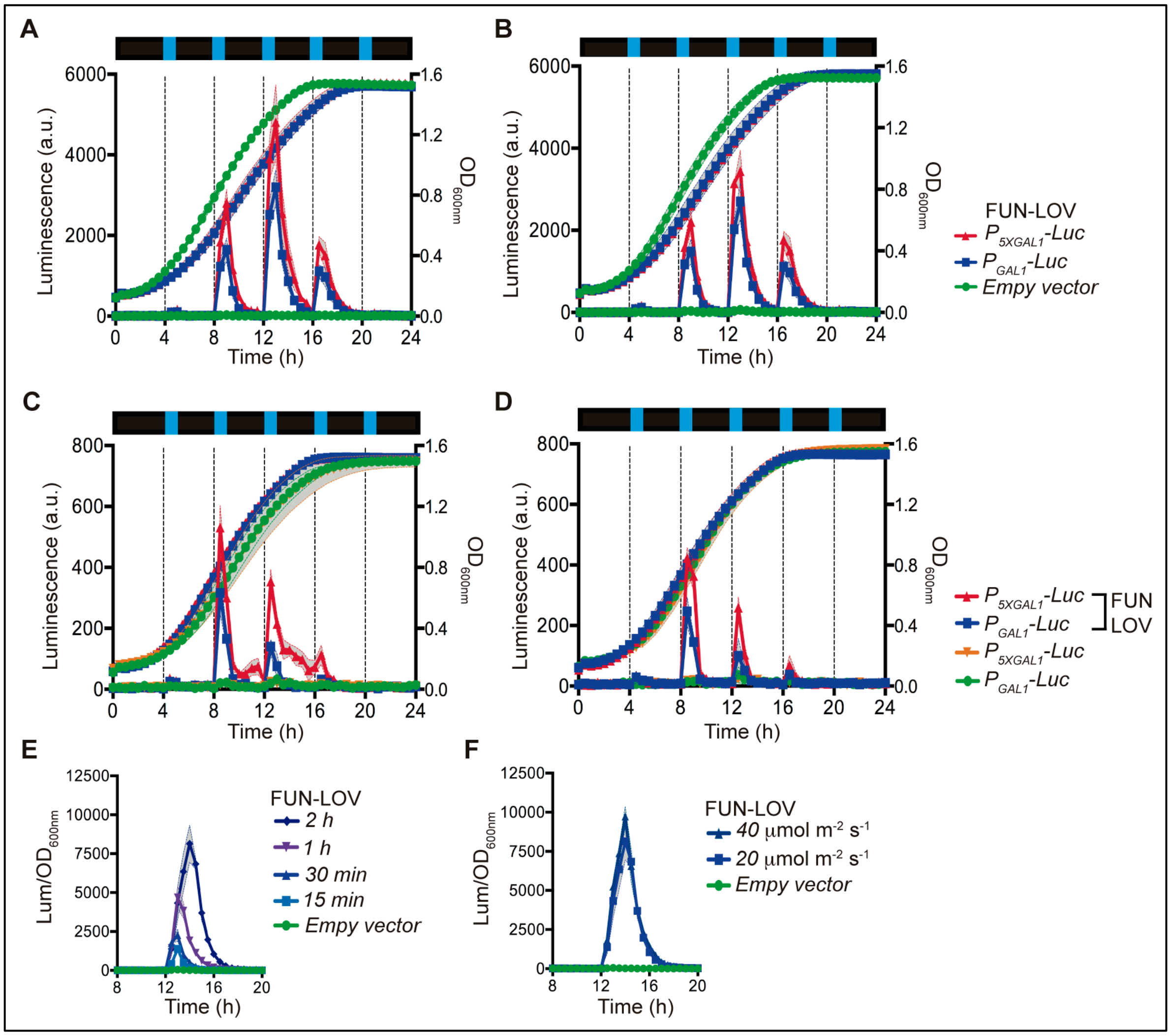


**FIG S3. Tuneable and dynamic control of luciferase expression by the FUN-LOV switch.** Bioluminescence was actively monitored as yeast cells were grown in constant darkness and 30-min blue-light pulses were applied as depicted (A-D). $P_{GAL1}$*-Luc* or $P_{5XGAL1}$*-Luc* reporters were maintained episomally in a BY4741 (A) or BY4741 *gal4*Δ*-gal80*Δ (B) background or inserted at the *GAL3* locus in BY4741 (C) or BY4741 *gal4*Δ*-gal80*Δ (D). The effect of the duration, or intensity of a blue-light pulse was evaluated in a strain containing an episomally maintained $P_{GAL1}$*–Luc* reporter (E and F respectively). In (F) the duration of the pulse set to 2 h. In panels (A) to (F), each plot corresponds to the average of six biological replicas with its standard deviation represented as shadowed regions.

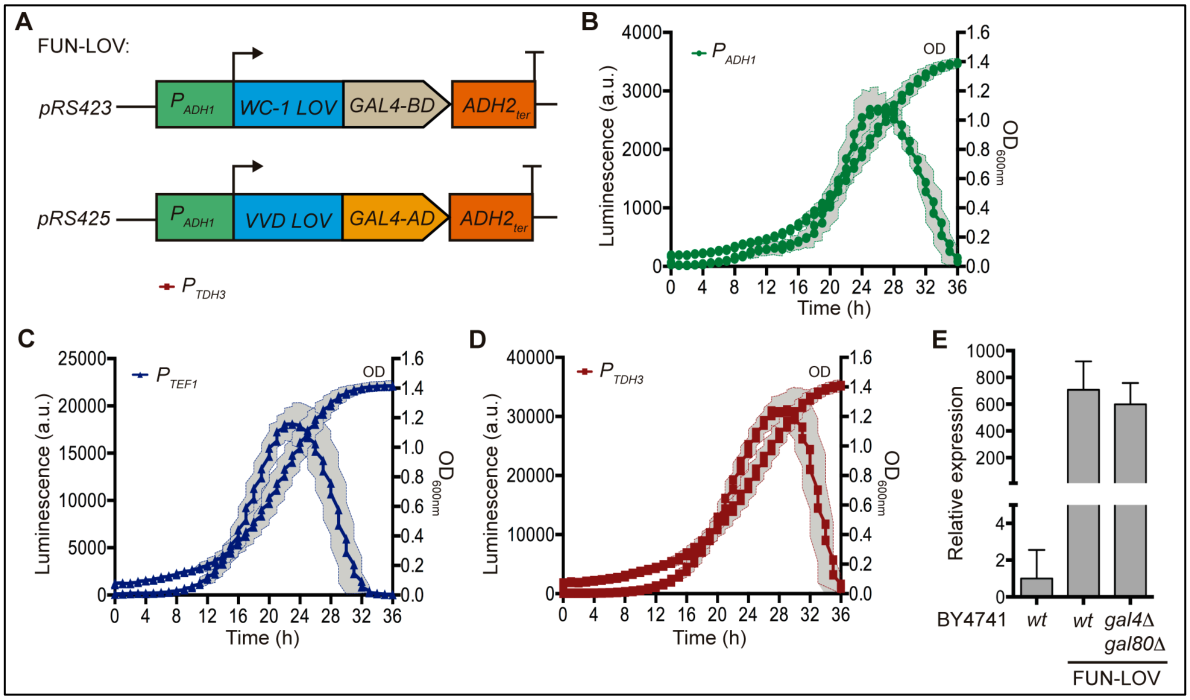


**FIG S4. Behaviour of the constitutive promoter controlling the FUN-LOV components.** (A) Expression of the FUN-LOV switch components (see also Fig 1A) is regulated by the *ADH1* promoter ($P_{ADH1}$) and *ADH2* transcriptional terminator (*ADH2*$_{ter}$). (B) Luciferase expression under the control of $P_{ADH1}$ promoter showed strong transcriptional activity during the exponential phase of the growth curve. (C-D) Luciferase expression under the control of two different constitutive promoters, $P_{TEF1}$ (C) and $P_{TDH3}$ (D), showing a similar behaviour to panel (B) but with higher expression levels, as revealed by the luminescence values. (E) The *GAL4* DNA binding domain (*GAL4-DBD*) linked to the WC-1 LOV domain is one of the components of the FUN-LOV system. *GAL4-DBD* expression was measured by real time PCR (qPCR) in two different genetic backgrounds (BY4741 and BY4741 *gal4Δ*/*gal80Δ*) and with/without the FUN-LOV system, confirming the strong expression of this component of the system. In panels (B) to (D), each plot corresponds to the average of three biological replicas with its standard deviation represented as shadowed regions. The qPCR experiments were performed in three biological replicas (n = 3), the average of these replicas with its standard deviation is shown in panel (E).

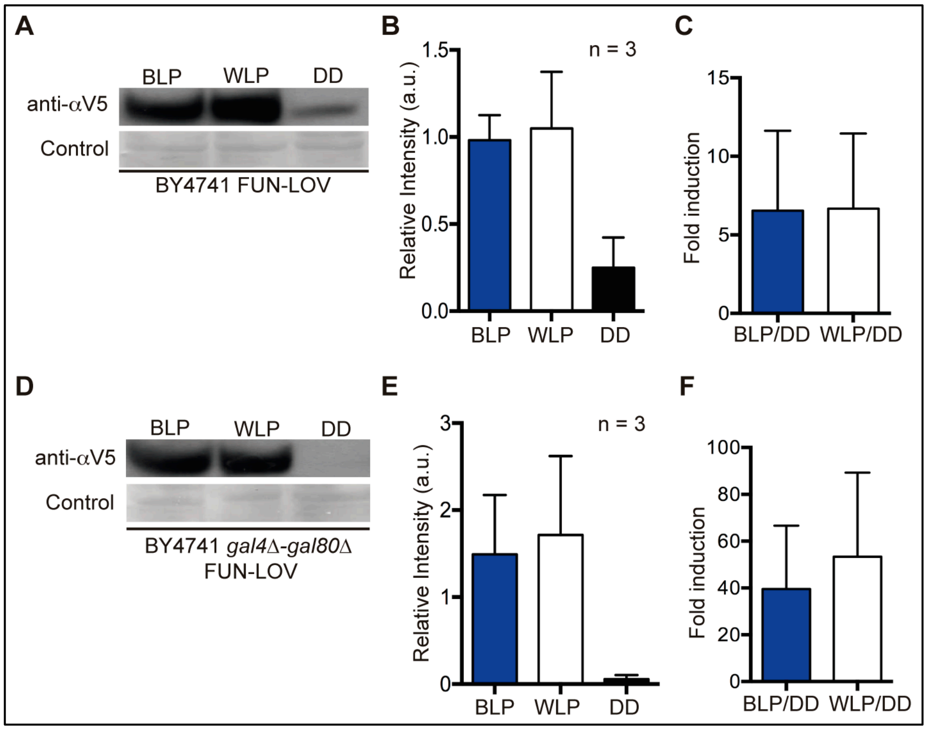


**FIG S5. FUN-LOV controlled expression of a heterologous protein.** (A-C) Expression of Limonene Synthase (LS) protein was analysed by Western blot (A) and quantified respect to membrane stained proteins (B) in three conditions: after 2 hours of blue-light pulse (BLP), after 2 hours of white-light pulse (WLP) and in constant darkness (DD). The fold of induction for LS expression was calculated after the BLP and WLP respect to DD (C). The experiments were carr ied out in the BY4741 wild type strain. (D-F) Similar experiments to panels (A-C) performed in the BY4741 *gal4Δ-gal80Δ* genetic background, which reduced the background expression in DD increasing the fold induction of the system.

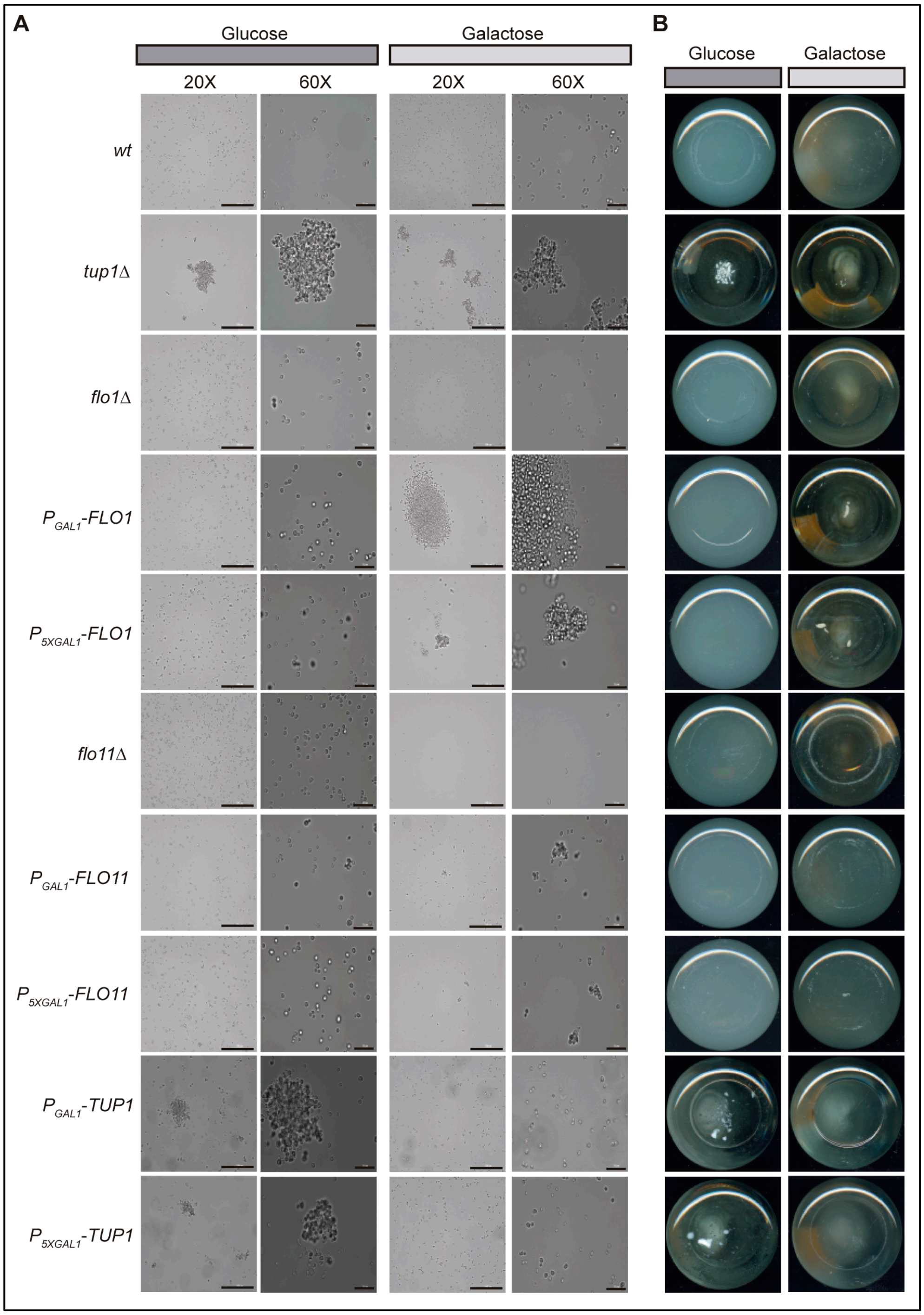


**FIG S6. Galactose and glucose control of the flocculation phenotype.** (A-B) Strains with *FLO1*, *FLO11* and *TUP1* promoter swapping ($P_{GAL1}$ and $P_{5XGAL1}$) were grown in galactose and glucose as carbon sources; and the flocculation phenotype was assayed by bright-field microscopy (A) and macroscopic analyses of the culture flasks (bottom view) (B). BY4741 and BY4741 *flo1*Δ strains were used as negative control of flocculation; whereas the BY4741 *tup1*Δ was used as positive control of flocculation. Pictures were taken with 20X and 60X objectives using a Cytation 3 in microscope mode, scale bar represent 100 μm at 20X and 20 μm at 60X, respectively.

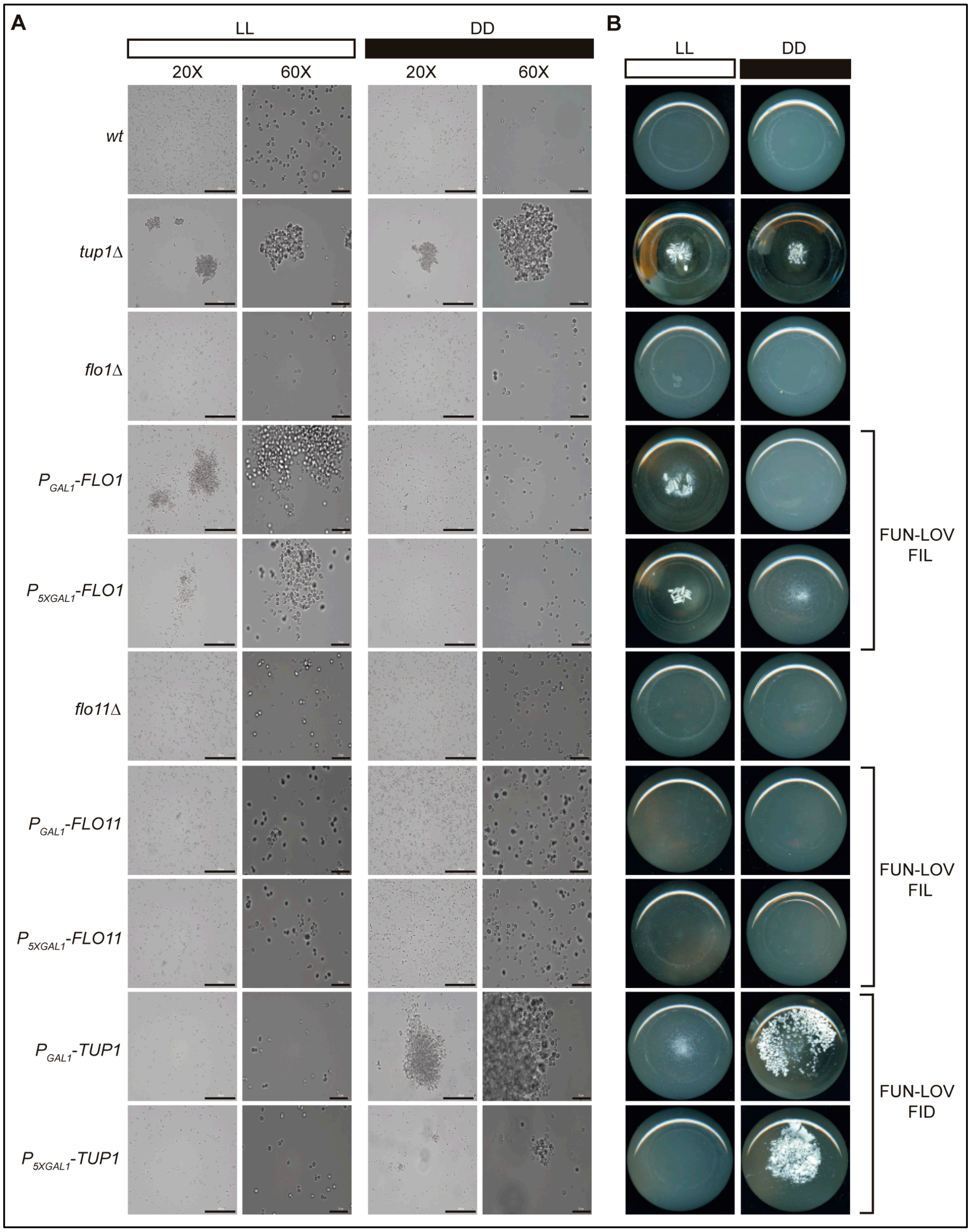


**FIG S7. FUN-LOV control of flocculation in light (FIL) and flocculation in Darkness (FID) phenotypes.** (A-B) Strains with *FLO1*, *FLO11* and *TUP1* promoter swapping ($P_{GAL1}$ and $P_{5XGAL1}$) and carrying the FUN-LOV system were grown under constant light (LL) and constant darkness (DD) conditions; and the flocculation phenotype was assayed by bright-field microscopy (A) and pictures of the culture flasks (bottom view) (B). The BY4741 *wt* and BY4741 *flo1*Δ strains were used as negative control of flocculation; whereas the BY4741 *tup1*Δ strain was used as positive control of flocculation. Pictures were taken with 20X and 60X objectives using a Cytation 3 in microscope mode, scale bar represent 100 µm at 20X and 20 µm at 60X, respectively.
