## Supplementary Materials for "FUN-LOV: Fungal LOV domains for optogenetic control of gene expression and flocculation in yeast"

**Table S1. Primers used in this work**

**For plasmids assembly utilizing yeast recombinational cloning.**

| Name | Sequence (5'-3') | Length (bp) | | Description |
|---|---|---|---|---|
| 13 | AGCGGATAACAATTTCACACAGGAA ACAGCTAGGCGCATGCAACTTCTTT | 50 | FW | *YRP-L-*$P_{ADH1}$ |
| oL3148 | GGTAACGCCAGGGTTTTCCCAGTCA CGACGCCGGTAGAGGTGTGGTCAAT | 50 | RV | $ADH2_{ter}$*-YRP-R* |
| 19 | AGCGGATAACAATTTCACACAGGAA ACAGCATCGATGAATTCGAGCTCGT | 50 | FW | *YRP-L-KanMxRV* |
| 20 | CGGGTTAATTAAGGCGCGCC | 20 | RV | *KanMxRV* |
| 21 | AAACAGATCTGGCGCGCCTTAATTA ACCCGACTAGTACGGATTAGAAGCC | 50 | FW | *KanMxRV-*$P_{GAL1}$ |
| oL3026 | aaacagatctggcgcgccttaatta acccgcggaggacagtactccgctc | 50 | FW | *KanMxRV-*$P_{5XGAL1}$ |
| oL3080 | GCCCTTCTTAATGTTCTTAGCATCG GCCATGGTAAGCTTAATATTCCCTA | 50 | RV | *Luc-*$P_{GAL1}$ |
| oL3079 | CGACTCACTATAGGGAATATTAAGC TTACCATGGCCGATGCTAAGAACAT | 50 | FW | $P_{GAL1}$*-Luc* |
| oL3081 | GGTAACGCCAGGGTTTTCCCAGTCA CGACGTGGATCCTTGCAAATTAAAG | 50 | RV | $CYC1_{ter}$*-YRP-R* |
| 22 | GGTAACGCCAGGGTTTTCCCAGTCA CGACGGGTAAGCTTAATATTCCCTA | 50 | RV | $P_{GAL1}(P_{5XGAL1})$*-YRP-R* |
| oL3150 | AGCGGATAACAATTTCACACAGGAA ACAGCATTTCAAAGAATACGTAAAT | 50 | FW | *YRP-L-*$P_{TDH3}$ |
| oL3151 | CATGTTATCCTCCTCGCCCTTGCTC ACCATTTTGTTTGTTTATGTGTGTT | 50 | RV | *mCherry-*$P_{TDH3}$ |
| oL3152 | GTTTCGAATAAACACACATAAACAA ACAAAATGGTGAGCAAGGGCGAGGA | 50 | FW | $P_{TDH3}$*-mCherry* |
| oL3153 | GAATGTAAGCGTGACATAACTAATT ACATGCTACTTGTACAGCTCGTCCA | 50 | RV | $CYC1_{ter}$*-mCherry* |
| oL3154 | ACCGGCGGCATGGACGAGCTGTACA AGTAGCATGTAATTAGTTATGTCAC | 50 | FW | *mCherry-*$CYC1_{ter}$ |
| oL3266 | TTTTGGGACGCTCGAAGGCTTTTAA TTTGCcgggttaattaaggcgcgcc | 50 | FW | $CYC1_{ter}$*-HphMx* |
| oL3267 | aaacagatctggcgcgccttaatta acccgGCAAATTAAAAGCCTTCGAG | 50 | RV | *HphMx-*$CYC1_{ter}$ |
| oL2562 | gGTAACGCCAGGGTTTTCCCAGTCA CGACGatcgatgaattcgagctcgt | 50 | RV | *HphMx-YRP-R* |

**Primers used for promoter swapping**

| Name | Sequence (5'-3') | Length (bp) | | Description |
|---|---|---|---|---|
| oL3024 | ctctttttcctaataaggtggagct tttggcttccagtatgctttcacgg atcgatgaattcgagctcgt | 70 | FW | $P_{FLO1\Delta}$::*KanMxRV-*$P_{GAL1}(P_{5XGAL1})$ |
| oL3025 | gccagaagtgtaaagactgccaaaa acatatagcgatgaggcattgtcat ggtaagcttaatattcccta | 70 | RV | $P_{FLO1\Delta}$::*KanMxRV-*$P_{GAL1}(P_{5XGAL1})$ |

| | | | | |
|---|---|---|---|---|
| oL3055 | ATTCTCATCGAGAGCCGAGCCATAC ACCTAAGGTGGACAGAAAGCTAAAA ATCGATGAATTCGAGCTCGT | 70 | FW | $P_{FLO11\Delta}$::KanMxRV-$P_{GAL1}$($P_{5XGAL1}$) |
| oL3056 | TTAAATAGAAGCGAAAGGACCAAAT AAGCGAGTAGAAATGGTCTTTGCAT GGTAAGCTTAATATTCCCTA | 70 | RV | $P_{FLO11\Delta}$::*KanMxRV*-$P_{GAL1}$($P_{5XGAL1}$) |
| oL3233 | CTTCACTCTTGACGCCAGTTGAGTG AGGGACTGAATACTATATACAGTAA ATCGATGAATTCGAGCTCGT | 70 | FW | $P_{TUP1\Delta}$::*KanMxRV*-$P_{GAL1}$($P_{5XGAL1}$) |
| oL3261 | TCGAGAAGCTCATTCAGCTTATTCT GCGTATTCGAAACGCTGGCAGTCATgg taagcttaatattcccta | 70 | RV | $P_{TUP1\Delta}$::*KanMxRV*-$P_{GAL1}$($P_{5XGAL1}$) |

**Primers used promoter-swapping confirmations.**

| Name | Sequence (5'-3') | Length (bp) | | Description |
|---|---|---|---|---|
| oL2789 | ttggtaacgatgagggaacc | 20 | FW | Upstream *FLO1* |
| oL3050 | ACTTTTCCTCTGGCCTGCTG | 20 | RV | Internal *FLO1* |
| oL3163 | TACCGTCGTTTTGGGGTTCC | 20 | FW | Upstream *FLO11* |
| oL3071 | TTGGGACAGCCATTAACGAT | 20 | RV | Internal *FLO11* |
| oL3262 | GCGGAATCGATCTGTTGTTT | 20 | FW | Upstream *TUP1* |
| oL3263 | CTTCTTCGTACGCGTCCTTC | 20 | RV | Internal *TUP1* |
| 21 | AAACAGATCTGGCGCGCCTTAATTA ACCCGACTAGTACGGATTAGAAGCC | 50 | FW | *KanMxRV*-$P_{GAL1}$ |
| oL3026 | aaacagatctggcgcgccttaatta acccgcggaggacagtactccgctc | 50 | FW | *KanMxRV*-$P_{5XGAL1}$ |
| oL2091 | CATCCTATGGAACTGCCTCGG | 21 | FW | Internal *KanMx* |

**Primers used for gene deletion and reporter gene integration in the genome.**

| Name | Sequence (5'-3') | Length (bp) | | Description |
|---|---|---|---|---|
| oL2787 | Ctctttttcctaataaggtggagct tttggcttccagtatgctttcacgg cggcatcagagcagattgtactg | 73 | FW | *flo1*Δ::*URA3* |
| oL2806 | Tactcgaaaacatcctaagcgaacc acactagatcttacgttagtactgc ggtatttcacaccgcataggg | 71 | RV | *flo1*Δ::*URA3* |
| oL3072 | TGCGGTATCTTCACGGACAGAACTT CTATTGCCTATCGGTGGTGTGATTA cggcatcagagcagattgtactg | 73 | FW | *flo11*Δ::*URA3* |
| oL3073 | CGAAGATTATTAGTTGTGCCAAGGC AATATCAGGTTTATTAATCTTTTAG ggtatttcacaccgcataggg | 71 | RV | *flo11*Δ::*URA3* |
| oL2120 | CCAGCGTATACAATCTCGATAGTTG GTTTCCCGTTCTTTCCACTCCCGTC | 70 | FW | *gal80*Δ::*HphMx* |

| | | | | |
|---|---|---|---|---|
| | CGGGTTAATTAAGGCGCGCC | | | |
| oL2121 | GTTTTTATAACGTTCGCTGCACTGG<br>GGGCCAAGCACAGGGCAAGATGCTT<br>ATCGATGAATTCGAGCTCGT | 70 | RV | *gal80Δ::HphMx* |
| oL2122 | ATGCACAGTTGAAGTGAACTTGCGG<br>GGTTTTTCAGTATCTACGATTCATT<br>CGGGTTAATTAAGGCGCGCC | 70 | FW | *gal4Δ::NatMx* |
| oL2123 | AATGCACGCCATCATTTTAAGAGAG<br>GACAGAGAAGCAAGCCTCCTGAAAG<br>ATCGATGAATTCGAGCTCGT | 70 | RV | *gal4Δ::NatMx* |
| oL3082 | AGGAGTGCAAAAAGAGAAAATAAAA<br>GTAAAAAGGTAGGGCAACACATAGT<br>atcgatgaattcgagctcgt | 70 | FW | *gal3Δ::KanMxRV-*<br>$P_{GAL1}$($P_{5XGAL1}$)*-Luc* |
| oL3083 | TATGAGTAAACTTTTAATATTTAAA<br>GGTTGTTCCAAGAAGGTGTTTAGTG<br>TGGATCCTTGCAAATTAAAG | 70 | RV | *gal3Δ::KanMxRV-*<br>$P_{GAL1}$($P_{5XGAL1}$)*-Luc* |

**Primers used for confirmation of gene deletions and genome integration.**

| Name | Sequence (5'-3') | Length (bp) | | Description |
|---|---|---|---|---|
| oL2789 | ttggtaacgatgagggaacc | 20 | FW | Upstream *FLO1* |
| oL2289 | CCTCTAGGTTCCTTTGTTACTTCT | 24 | RV | Internal *URA3* |
| oL2807 | tcacctgctttgaatcgttg | 20 | RV | Upstream *FLO1* |
| oL2288 | CCTTTTGATGTTAGCAGAATTGTC | 24 | FW | Internal *URA3* |
| oL2099 | GAGACAGCATTCGCCCAGTA | 20 | FW | Upstream *GAL4* |
| oL2163 | GATTCGTCGTCCGATTCGTC | 20 | RV | Internal *NatMx* |
| 4 | AGGTTACATGGCCAAGATTGA | 21 | RV | Upstream *GAL4* |
| oL2164 | AGGTCACCAACGTCAACGCA | 20 | FW | Internal *NatMx* |
| oL2096 | TCTTCATTTACCGGCGCACT | 20 | FW | Upstream *GAL80* |
| oL2094 | CGGCGGGAGATGCAATAGG | 19 | RV | Internal *HphMx* |
| oL2097 | CGCTGCTGCAAAGTTTTGAC | 20 | RV | Upstream *GAL80* |
| oL2095 | TCGCCCGCAGAAGCGCGGCC | 20 | FW | Internal *HphMx* |
| oL3078 | CAGTCCGAGCGTTTAGAAGG | 20 | FW | Upstream *FLO11* |
| oL3089 | ATGAAATCGCCATGCCAAGC | 20 | FW | Upstream *GAL3* |
| oL2091 | CATCCTATGGAACTGCCTCGG | 21 | FW | Internal *KanMx* |
| oL3127 | GTGCGGAGCCACTCTGACTC | 20 | RV | Upstream *GAL3* |
| oL1734 | GGCCGGCCGCTTCGAGCAGACATGA<br>TAAGAtcatgtaattagttatgtca | 50 | FW | $CYC1_{ter}$*-Luc* |

**Primers used for real time PCR (qPCR).**

| Name | Sequence (5'-3') | Length (bp) | | Description |
|---|---|---|---|---|
| oL3682 | GGGTTTGCGTTCTGTGATCT | 20 | FW | *HEM2 housekeeping* |
| oL3683 | TTGAATGACTGGCCCTGCAG | 20 | RV | *HEM2 housekeeping* |
| oL3692 | GGATCAGGTCTTCCGTAGCG | 20 | FW | *TAF10 housekeeping* |
| oL3693 | TGAAATCTGCTGCACGCCA | 19 | RV | *TAF10 housekeeping* |
| oL3321 | TGCCGTCACAGATAGATTGG | 20 | RV | *GAL4-BD* |
| oL3322 | CTCTTCCGATGATGATGTCG | 20 | FW | *GAL4-BD* |
